# Delivery of A Jagged1-PEG-MAL hydrogel with Pediatric Human Bone Cells Regenerates Critically-Sized Craniofacial Bone Defects

**DOI:** 10.1101/2023.10.06.561291

**Authors:** Archana Kamalakar, Brendan Tobin, Sundus Kaimari, M. Hope Robinson, Afra I. Toma, Timothy Cha, Samir Chihab, Irica Moriarity, Surabhi Gautam, Pallavi Bhattaram, Shelly Abramowicz, Hicham Drissi, Andrés J. García, Levi B. Wood, Steven L. Goudy

**Affiliations:** Department of Pediatric Otolaryngology, Emory University, Atlanta, GA, USA; Department of Cell biology, Emory University, Atlanta, GA, USA; Department of Orthopedics, Emory University, Atlanta, GA, USA; Department of Surgery, Division of Oral and Maxillofacial Surgery, Emory University, Atlanta, GA, USA; Department of Pediatric Otolaryngology, Children’s Healthcare of Atlanta, Atlanta, GA, USA; Wallace H. Coulter Department of Biomedical Engineering, Georgia Institute of Technology, Atlanta, GA, USA; Neuroscience Program in College of Sciences, Georgia Institute of Technology, Atlanta, GA, USA; Parker H. Petit Institute for Bioengineering and Biosciences, Georgia Institute of Technology, Atlanta, GA, USA; George W. Woodruff School of Mechanical Engineering, Georgia Tech College of Engineering, Atlanta, GA, USA; The Atlanta Veterans Affairs Medical Center Atlanta, GA, USA; School of Chemistry and Biomolecular Engineering, Georgia Tech College of Engineering, Atlanta, GA, USA

**Keywords:** JAGGED1, Human Bone-derived Osteoblast-like Cells, PEG-4MAL Hydrogel-based delivery, Craniofacial Bone Defects, Bone Regeneration, Non-canonical JAG1-NOTCH pathways, RNA Sequencing

## Abstract

Treatments for congenital and acquired craniofacial (CF) bone abnormalities are limited and expensive. Current reconstructive methods include surgical correction of injuries, short-term bone stabilization, and long-term use of bone grafting solutions, including implantation of (i) allografts which are prone to implant failure or infection, (ii) autografts which are limited in supply. Current bone regenerative approaches have consistently relied on BMP2 application with or without addition of stem cells. BMP2 treatment can lead to severe bony overgrowth or uncontrolled inflammation, which can accelerate further bone loss. Bone marrow-derived mesenchymal stem cell-based treatments, which do not have the side effects of BMP2, are not currently FDA approved, and are time and resource intensive. There is a critical need for novel bone regenerative therapies to treat CF bone loss that have minimal side effects, are easily available, and are affordable. In this study we investigated novel bone regenerative therapies downstream of JAGGED1 (JAG1).

We previously demonstrated that JAG1 induces murine cranial neural crest (CNC) cells towards osteoblast commitment via a NOTCH non-canonical pathway involving JAK2-STAT5 (1) and that JAG1 delivery with CNC cells elicits bone regeneration in vivo. In this study, we hypothesize that delivery of JAG1 and induction of its downstream NOTCH non-canonical signaling in **pediatric human osteoblasts** constitute an effective bone regenerative treatment in an in vivo murine bone loss model of a critically-sized cranial defect. Using this CF defect model in vivo, we delivered JAG1 with pediatric human bone-derived osteoblast-like (HBO) cells to demonstrate the osteo-inductive properties of JAG1 in human cells and in vitro we utilized the HBO cells to identify the downstream non-canonical JAG1 signaling intermediates as effective bone regenerative treatments. In vitro, we identified an important mechanism by which JAG1 induces pediatric osteoblast commitment and bone formation involving the phosphorylation of p70 S6K. This discovery enables potential new treatment avenues involving the delivery of tethered JAG1 and the downstream activators of p70 S6K as powerful bone regenerative therapies in pediatric CF bone loss.

## 1. Introduction

Craniofacial (CF) injuries comprise more than 25% of injuries reported to the National Trauma Data Bank in the US every year (2–4). Left untreated, CF injuries can severely impair critical daily functions related to breathing, speech, eating, and swallowing, thus requiring urgent repair (5). Current methods to repair CF bone loss include the direct implantation of either allografts or autografts, and/or subsequent revision surgeries (6, 7). Although generally successful, these conventional treatments present several limitations. Bone donor sites include rib, fibula, iliac crest, scapula, distal tibia, and medial femoral condyle. All these donor sites have limited availability and are not anatomically like the bone they replace; osteotomies are required to shape them so that they resemble the shape of the CF bones (8, 9). The allograft survival rate following an iliac graft procedure is 96.1% and the risk of infection ranges from 5 to 33% (7, 10). For maxillary or mandibular bone replacement, the iliac crest is the preferred graft source due to its robust corticocancellous anatomy. Risks associated with iliac crest grafting include significant pain at the donor site, nerve injury, decreased load bearing on the ipsilateral leg, and increased risk of hip fractures (11, 12). Due to the limited supply of bone, revision bone graft surgeries are often required and are expensive ($35,000 – $52,000 per patient) and cause great discomfort (13). These and other complications undermine the patient’s quality of life in addition to the social stigma resulting from the facial deformity, which can also lead to psychological distress (14).

Craniofacial bone development primarily occurs through intramembranous ossification, a process where pre-osteoblasts mineralize directly, without a cartilage intermediate, making it distinct from long bone development (endochondral ossification) (15). Intramembranous ossification recruits Cranial Neural Crest (CNC) cells as osteoblast precursors during CF bone development (15). Extensive efforts have been made towards using Parathyroid Hormone (PTH (1–34)), Vascular Endothelial Growth Factor (VEGF), Fibroblast Growth Factor (FGF), Stromal Cell-Derived Factor-1 (SDF-1) or Transforming Growth Factor- Beta 2 (TGFβ2) as alternatives to bone regenerative treatments in preclinical models but resulted in limited success as an in vivo treatment option (16–20). Platelet-rich Plasma (PRP) supplemented with various biological growth factors such as VEGF, FGF, and TGFβ2 have also been used to enhance wound healing, chemotaxis, angiogenesis, proliferation of mesenchymal stem cells and osteoblasts. Many of these studies demonstrated potential improvement in bone healing; however, these strategies face significant translational barriers (20). Bone Morphogenetic Protein-2 (BMP2) is an FDA-approved bone regenerative strategy along with the delivery of stem cells. BMP2 is used to reconstruct spine and maxillary bone in adults (21, 22). BMP2 treatment can lead to ectopic bone growth, hypertrophy and life-threatening side effects (e.g., uncontrolled inflammation), which may accelerate bone loss (23, 24). The use of BMP2 for the treatment of pediatric cases of CF trauma is not FDA-approved due to concerns of severe swelling of the face and airways. Although generally considered safer, stem cell-based treatments are time consuming, have heterogenous results and add to the high expense of repairing CF bone loss (22). *Thus, there is a critical need for novel bone regenerative therapies to treat CF bone loss that have minimal side effects, are readily accessible and affordable*.

The NOTCH signaling pathway is involved in many cellular processes, including determination of cell fate, and has been explored as a potential target for regeneration of long bone injuries (25). NOTCH signaling occurs via cell-to-cell binding of a NOTCH ligand (e.g., JAG1) to a NOTCH receptor. Internalization of the NOTCH intracellular domain leads to the expression of canonical NOTCH genes *HES1* and *HEY1*, which are known to have both osteo-inductive and osteo-inhibitory roles, thus obfuscating the effectiveness of NOTCH-based bone regenerative therapies (26, 27). However, we and others have demonstrated that JAG1 exhibits osteo-inductive properties, as demonstrated by induction of pre-osteoblast genes like *Runx2* in murine CNC cells, even when the canonical NOTCH pathway is inhibited (1, 28). Our recent publications further identified a JAG1-JAK2 non-canonical signaling pathway that promotes murine CNC commitment to osteoblast differentiation in mice (1).

This unexpected finding raises critical questions about the role of non-canonical JAG1 signaling during human craniofacial regeneration. ***In this study we postulate that 1) JAG1 can induce osteoblast differentiation and mineralization of pediatric Human Bone-derived Osteoblast-like (HBO) cells, and 2) the delivery of JAG1 non-canonical signaling constitutes an effective treatment for inducing bone regeneration in a pediatric, preclinical craniofacial bone loss model.*** Consequently, we evaluated (i) the ability of JAG1 to regenerate bone in a pediatric critically-sized CF defect murine model when delivered in a synthetic hydrogel coupled with pediatric HBO cells, and (ii) characterized the downstream JAGGED1 non-canonical signaling mechanisms. Our results reveal potential treatment options in the form of JAG1 and/or its downstream targets to induce bone regeneration in CF bone loss injuries and provide alternative treatment options for CF defects.

## 2. Materials & Methods

### 2.1 HBO cell isolation

HBO cell lines were derived from healthy fibulas of seven pediatric subjects under appropriate Institutional Review Board approval. Segments of the fibulas were digested using collagenase A (Roche, 10103578001) used at 0.1 mg/mL treatment for 40 minutes with replacement of collagenase A midway at 20 minutes and then digested with 0.2 mg/ml of collagenase A for 60 minutes at 37°C with intermittent shaking.

The segments of bone were maintained in DMEM + Primocin® Antimicrobial agent for primary cells (Invivogen, ant-pm-1) + 10% FBS + 50 µM ascorbic acid (Sigma, 49752) + 10nM dexamethasone. Media changes were performed every 5 days. Osteoblast-like cells began to grow out of the bone segments by Day 10. These cells can be passaged and frozen for storage. HBO cell lines used for experiments were selected based on the ability of the primary cell line to proliferate and mineralize in culture. On addition of Osteogenic media [(DMEM + Primocin + 10% FBS + 50 µM ascorbic acid + 10 nM dexamethasone (Sigma, D1756) + 10 mM beta-glycerophosphate disodium (Sigma, G9422)], the cells form mineralized nodules indicating their osteogenic ability as seen in **Supplementary Figure 1**.

### 2.2 JAG1 immobilization

As per manufacturer’s recommendations, 50 μL (1.5 mg) of Dynabeads Protein G (Invitrogen 10004D) were transferred to a tube, where the beads were separated from the solution using a magnetic tube rack (Biorad, 1614916) and washed once using 200 μL PBS with 0.1% Tween-20 (Fisher, BP337-500). The wash buffer was separated from the beads-Fc complex using the magnetic rack. Recombinant JAG1-Fc (5 µM, 5.7 µM, 10 µM or 20 µM) (Creative Biomart, JAG1-3138H) or control IgG-Fc fragment (5 µM or 5.7 µM) (Abcam, ab90285) were diluted in 200 μL PBS with 0.1% Tween-20 and then added to the Dynabeads. The beads plus proteins were incubated at 4°C with rotation for 16 hours. Thereafter, the tubes were placed back on the magnetic rack and the supernatant was removed. The bead-JAG1/Fc complex was resuspended in 200 μL PBS with 0.1% Tween-20 to wash by gentle pipetting. The wash buffer was also separated from the beads-Fc complex using the magnetic rack, and the final suspension of the beads in hydrogels was used as treatment.

### 2.3 Mineralization assay

HBO cells were seeded at 30,000 cells per well in a 12-well plate, treated (n = 3) with and cultured for 21 days in osteogenic media (DMEM + Primocin + 10% FBS + 50 µM ascorbic acid + 10 nM dexamethasone + 10 mM beta-glycerophosphate) with half feeds every 5 days. Treatments included growth media (DMEM + Primocin + 10% FBS), osteogenic media, Fc-Dynabeads (5.7 μM) in osteogenic media or JAG1-Dynabeads (5.7 μM) in osteogenic media. Inhibitors N-[N-(3,5-Difluorophenacetyl)-L-alanyl]-S-phenylglycine t-butyl ester (DAPT; Sigma, D5942) (15 µM) and 5-(1,1-dimethylethyl)-2-[[(1H-indazol-5- ylamino)carbonyl]amino]-3-thiophenecarboxylic acid (S6K-18 – inhibitor of p70 ribosomal S6 kinase 1; Selleck Chemicals, S0385) (50 μM) were also added to JAG1-Dynabeads conditions for experiments shown in Figure 5 and Supplementary Figures 6 and 7. For PCR, cells were collected on days 7-9, 14, and 21 (see Methods 2.8.1).

For Alizarin Red staining: On day 21, the cells were fixed using 50% ethanol for 15 minutes at 4°C. The fixed cells were then stained with Alizarin Red S dye (LabChem, LC106002) to detect mineralization. The dye was extracted using a 10% acetic acid solution in water and quantified by measuring the absorbance at 420nm using a spectrophotometer.

### 2.4 Hydrogel preparation

We prepared poly (ethylene glycol) (PEG)-based synthetic hydrogels incorporating cell adhesive peptides in two steps. First, maleimide end-functionalized 20 kDa four-arm PEG macromer (PEG-4MAL, with > 95% end-group substitution, Laysan Bio, 4ARM-PEG-MAL-20K), was reacted with a thiol-containing adhesive peptide GRGDSPC (Genscript, RP20283) in PBS with 20 mM HEPES at pH 7.4 for 1 hour. Then, the RGD-functionalized PEG-4MAL macromers were cross-linked in the presence of HBO cells and JAG1-Dynabeads into a hydrogel by addition of the dithiol protease-cleavable peptide cross-linker GPQ-W (GCRDGPQGIWGQDRCG) (New England Peptides, Inc, (NEP) Custom synthesized) (1, 29). The final gel formulation consisted of 4.0% wt/vol polymer and 1.0 mM RGD.

### 2.5 In vivo experiments

We performed all in vivo experiments using procedural guidelines with appropriate approvals from the Institutional Animal Care and Use Committee of Emory University. 6-8-week-old male and female NOD-SCID mice (The Jackson Laboratory, 001303) were used. As shown in **Supplementary Figure 3**, the surgery site was disinfected and then incisions were made using sterile surgical equipment to expose the parietal bones of the mice. Thereafter, 4 mm defects were created in the parietal bones using a variable speed drill (Aseptico (MicroNX), MAX-88ESP, CL1791023) and sterile circular knives. **JAG1 Delivery:** PEG-4MAL hydrogels (20 µL) loaded with JAG1-Dynabeads without or with 100,000 HBO cells (n = 13 – 15 for all treatment groups) were placed within the defects created in parietal bones in the NOD-SCID mouse skulls as the first dose. A second dose of the hydrogels encapsulating HBO cells and Dynabead-bound JAG1 were administered as transcutaneous injections during week four to continue the bone regenerative action of JAG1. Skulls were then harvested at week eight, fixed using 10% neutral buffered formalin (VWR, 89370-094) and imaged with micro computed tomography (µCT). HBO primary cell lines 2, 6 and 7 from separate individuals were selected for these experiments based on similar growth and passage characteristics.

### 2.6 Micro Computed Tomography (µCT)

µCT analyses were conducted according to current guidelines for the assessment of bone volume within the defects created in mouse calvaria (30). Briefly, formalin-fixed skulls were positioned in the µCT tubes with the nose facing the bottom of the tube and imaged in a µCT 40 (Scanco Medical AG, Bassersdorf, Switzerland) using a 36 µm isotropic voxel size in all dimensions.

Thereafter, using a consistent and pre-determined threshold of 55 kVp, 145 µA, 8 W and 200 ms integration time for all measurements, three-dimensional (3D) reconstructions were created by stacking the regions of interest from ∼600 two-dimensional (2D) slices consisting of the entire skull and then applying a gray-scale threshold of 150 and Gaussian noise filter (σ= 0.8, support = 1.0), a coronal reformatting was done. Thereafter, a circular region of interest (ROI) encompassing the defect was selected for analysis consisting of transverse CT slices encompassing the entire defect and, new bone volume (BV) was calculated.

### 2.7 Histology

#### 2.7.1 Masson Trichrome staining

Formalin-fixed skulls used for µCT measurements were decalcified in Cal- ex (Fisher, C5510-1D), embedded in paraffin, and sectioned on a microtome to obtain 5 µm sections which were then stained using a Masson Trichrome staining kit (Sigma Aldrich, HT15) according to the manufacturer’s protocol. The sections were first washed with PBS, three times, for 5 minutes each, then the slides were submerged in Bouin’s solution for 15 minutes, and then washed under running water for 5 minutes. Thereafter, the sections were incubated in Weigert’s working hematoxylin solution for 10 minutes before being washed three times under running water for 5 minutes each and then transferred to distilled water. The sections were then stained with biebrich scarlet acid fuchsin for 5 minutes and washed with distilled water three times before being submerged in a phosphotungstic/phosphomolybdic solution for 10 minutes, and subsequently placed in an analine blue solution. Lastly, they were washed with distilled water three times and moved to a solution of 1% acetic acid for 1 minute followed by a submersion in two changes of xylene and mounted with a coverslip. The sections were then imaged using brightfield microscopy.

#### 2.7.2 Immunohistochemical analysis

Deparaffinization of FFPE sections was first performed by incubating them for 45 minutes at 60°C followed by Xylene (2 times, 5 minutes each) and then sections were rehydrated using various grades of alcohol in a decreasing concentration (100%, 90% and 75%). The sections were washed with 1X PBS for 10 minutes, permeabilized with 1.0% Triton X-100 (in 1X PBS) for 10 minutes at room temperature and rinsed with 1X PBS (3 times, 5 minutes each). The sections were covered with 20 µg/ml of Proteinase K in 1X PBS for 20 minutes at 37°C in a humidified chamber. After a 1X PBS rinse (2 times, 5 minutes each), the sections were blocked with 1% Bovine Serum Albumin (Fraction V, BP1600-100) for 30 minutes at room temperature. The sections were then washed with 1X PBS (2 times, 5 minutes each) and incubated with the primary antibody (COL1A1, Cell signaling #72026S) at 1:50 dilution for 24h at 4°C. Recombinant rabbit IgG monoclonal antibody was used as a negative isotype control. Sections were washed with 1X PBS (3 times, 5 minutes each), covered with two drops of SignalStain Boost Detection Reagent (HRP, Rabbit #8114), and incubated in a humidified chamber for 30 minutes at room temperature. After 1X PBS wash (3 times, 5 minutes), SignalStain DAB Substrate (Kit #8059) was applied for 5 minutes. The slides were immersed in de-ionized water (dH2O) for 5 minutes, and then counterstained with Hematoxylin [(Vintage Hematoxylin (SL 100)] for 5 seconds. After a 5-minute wash with dH2O, the sections were dehydrated with alcohol in increasing concentration (95% and 100%) followed by two incubations in Xylene for 10 seconds each. The slides were mounted and coverslipped using Permount Mounting Medium (#17986-05). Slides were scanned with the Olympus Nanozoomer whole-slide scanner at 20x.

### 2.8 RNA extraction, qRT–PCR, and RNA-sequencing

#### 2.8.1 RNA extraction and qRT-PCR

HBO cells were grown in biological triplicates in growth media alone, osteogenic media alone or with Fc-Dynabeads (5.7 μM) or JAG1-Dynabeads (5.7 μM). Cells were collected at 7-9 days, 14 days, and 21 days. RNA was extracted using TRIzol™ (Invitrogen, 15596026) by adding the TRIzol (400µL– 1mL) directly to plates of adherent cells and collecting by pipette. Samples were frozen at - 80°C and then processed in batches. Samples were allowed to thaw, and the appropriate volume of chloroform was added (200µL per mL of TRIzol), samples were vortexed for 1 minute, allowed to rest for 5 minutes and then centrifuged at 12,000xg, 15 minutes, 6°C. After centrifugation, the aqueous phase was removed to a separate tube and cold isopropanol was added (600µL isopropanol per mL TRIzol). Samples were mixed well and kept at -20°C for 10 minutes followed by centrifugation at 12,000xg, 10 minutes, 6°C. Supernatant was removed and cold 75% ethanol/25% RNase free water was added (500µL per mL Trizol). Samples were vortexed and centrifuged at 12,000xg, 5 minutes, 6°C. Supernatant was removed and samples were allowed to air dry for 10 minutes. RNA was resuspended in 20-30µL of RNase free water, warmed at 55°C for 10 minutes, and then quantified using a NanoDrop One spectrophotometer. The High Capacity cDNA Reverse Transcription Kit (Ref 4368814) from Applied Biosystems was used to produce cDNA following the manufacturer’s instructions. Primers were acquired from Integrated DNA Technologies (see Supplementary

Table 1 for sequences) and qRT-PCR was done in duplicate using BioRad iQ SYBR Green Supermix (BioRad, 1708882) following manufacturer’s instructions. Data was normalized to growth media condition with GAPDH as the reference gene. Analysis of qRT-PCR was accomplished by using double delta Ct analysis in Excel and data was plotted in GraphPad Prism 10.

#### 2.8.2 RNA Sequencing

RNA was isolated using the Qiagen RNeasy kit (Qiagen, 74106) according to manufacturer’s protocols. The samples were submitted to the Molecular Evolution core at the Georgia Institute of Technology for sequencing. Quality Control (QC) was performed using an Agilent Bioanalyzer 2100 to determine the RNA Integrity Number (RIN) of the samples. mRNA was enriched using the New England Biolab’s (NEB) NEBNext Poly(A) mRNA isolation module for samples with RINs greater than 7 and libraries were prepared using the NEBNext Ultra II directional RNA library preparation kit (NEB, E7760). QC was then performed on these libraries using an Agilent Bioanalyzer 2100 and the libraries were quantified using fluorometric methods. Paired-end 150 base pairs (PE150) sequencing was performed on the Illumina NovaSeq 6000 instrument to obtain a sequencing depth of 30 million reads per sample. The transcripts obtained were aligned using the hg38 genome reference database along with elimination of duplicate reads, using the DNAStar Lasergene 17.3 application. The RNA levels were calculated in reads per kilobase per million mapped reads (RPKM). Genes expressed at > 1.5 RPKM were retained for further analyses.

### 2.9 Transcriptomic Analysis Methods

#### 2.9.1 Differential Gene Expression and Enrichment Analysis

Differentially expressed genes (DEGs) were determined using DESeq2 (v1.38.3) available in R Bioconductor (31). Transcripts with an FDR-adjusted p-value<0.05 were considered significant for this analysis. Transcript counts were normalized using the median-of-ratios method used by DESeq2 prior to differential expression analysis. Results were visualized with Venn diagrams using govenn (v0.1.10), volcano plots in R using ggplot2 (v3.4.1) (H. Wickham. Ggplot2: Elegant Graphics for Data Analysis. Springer-Verlag New York, 2016.) and heatmaps in R using Heatmap3 (v1.1.9) (32).

#### 2.9.2 Transcript Functional Annotation

To provide additional functional annotation of the DEGs, a web-scraper was built to parse the National Center for Biotechnology Information Gene Database (33) entry for each transcript and identify relevant terms. Transcript names were connected to the NCBI Entrez ID with the Genome wide annotation for Human package, org.Hs.eg.db (v3.15.0), available through Bioconductor in R Carlson M (2019). *Org.Hs.eg.db: Genome wide annotation for Human*. R package version 3.15.0. This tool utilized the HTML parser in the BeautifulSoup4 (v4.12.2) Python package to extract the gene summary information from each gene entry in the database. The script then identified the keywords in the summary.

#### 2.9.3 Functional Over-Representation Analysis

Over-represented Gene Ontology (GO) terms were identified using PANTHER (v17.0) (34, 35). DEGs involved in the noncanonical signaling pathway were identified from Venn diagrams and separated as up-regulated or down-regulated relative to control. Each list was uploaded to the PANTHER online tool for over-representation testing. FDR-adjusted Fischer’s test p<0.05 was considered over-represented in this analysis. The complete GO Biological Process term list was evaluated and the complete list of identified transcripts from RNAseq was set as the background.

### 2.10 Luminex-based Multiplex assay

To detect phosphor-signaling targets, serum-starved HBO cell lines (n = 3) from different patients were treated (n = 3 per cell line) with Fc-Dynabeads (5.7 µM), unbound BMP2 (100 nM), and JAG1-dynabeads (5.7 µM) with or without DAPT (15 µM) as a time course stimulation for 5, 10, 15, and 30 minutes. Whole cell protein (2 µg) lysates were subjected to a Millipore Luminex based Multiplex assay to measure signaling targets using the Milliplex multiple pathway cell signaling magnetic bead 9-Plex kit (Millipore Sigma, 48-681MAG) according to manufacturer’s protocol.

### 2.11 Statistics

Data were analyzed by analysis of variance (ANOVA) with Tukey’s post-hoc test unless otherwise noted using GraphPad Prism 10. All data are presented as mean ± SD. P < 0.05 between groups was considered significant and are reported as such. MATLAB coding was used to create heatmaps and to generate z-score values associated with color intensities seen on the heatmap.

## 3. Results

### JAG1 induces mineralization of pediatric human bone derived osteoblast-like cells

To test whether JAG1 induces osteoblast commitment and differentiation in human osteoblast-like primary cells, we derived human bone osteoblast-like (HBO) cell lines (HBO1, 2, 3, 4, 5, 6 and 7) from seven healthy pediatric human bone samples, as described (36) (**Supplementary Figure 1**). HBO cells were then treated with Fc-Dynabeads (Fc-bds) (5.7 μM) as a negative control, since JAG1 is a chimeric recombinant protein with an Fc-portion, and JAG1-Dynabeads (JAG1-bds) (5.7 μM), in the presence of osteogenic media. On day 21, the cells were stained for mineralization using Alizarin Red S stain (**Figure 1A & 1B**). We observed significantly increased mineralization in JAG1-bds-treated samples compared to growth media-treated, osteogenic media-treated, and Fc-bds-treated (p = 0.0107) cells. Additionally, PCR analysis of HBO1 cells from a repeat experiment collected at days 7, 14, and 21 showed significantly increased expression of osteogenic genes with JAG1-bds stimulation (**Figure 1C**). *ALPL* was significantly expressed at Day 7, with a 3.5-fold increase (p=0.0004) compared to HBO1 cells grown in growth media. In contrast, significant expression levels of *COL1A1* and *BGLAP* were observed at 14 days, with a 5.1-fold increase (p=0.0021) of *COL1A1* and a 12.3-fold increase (0.0002) of *BGLAP* when compared to growth media conditions. Interestingly, while some mineralization is observed in the osteogenic media and Fc-bds **(Figure 1A)** conditions, there were no significant increases in osteogenic gene expression (**Figure 1C**). Expression of *RUNX2* and *SP7* was not significantly altered across all conditions and time points (not shown). In preparation for in vivo studies, HBO cells were incorporated into JAG1-Dynabead-PEG-4MAL and grown in vitro followed by Alizarin Red S staining to assess mineralization (**Supplementary Figure 2**). These data indicate that JAG1 can induce osteoblast commitment, differentiation, and mineralization of pediatric HBO cells.

**Figure 1:**
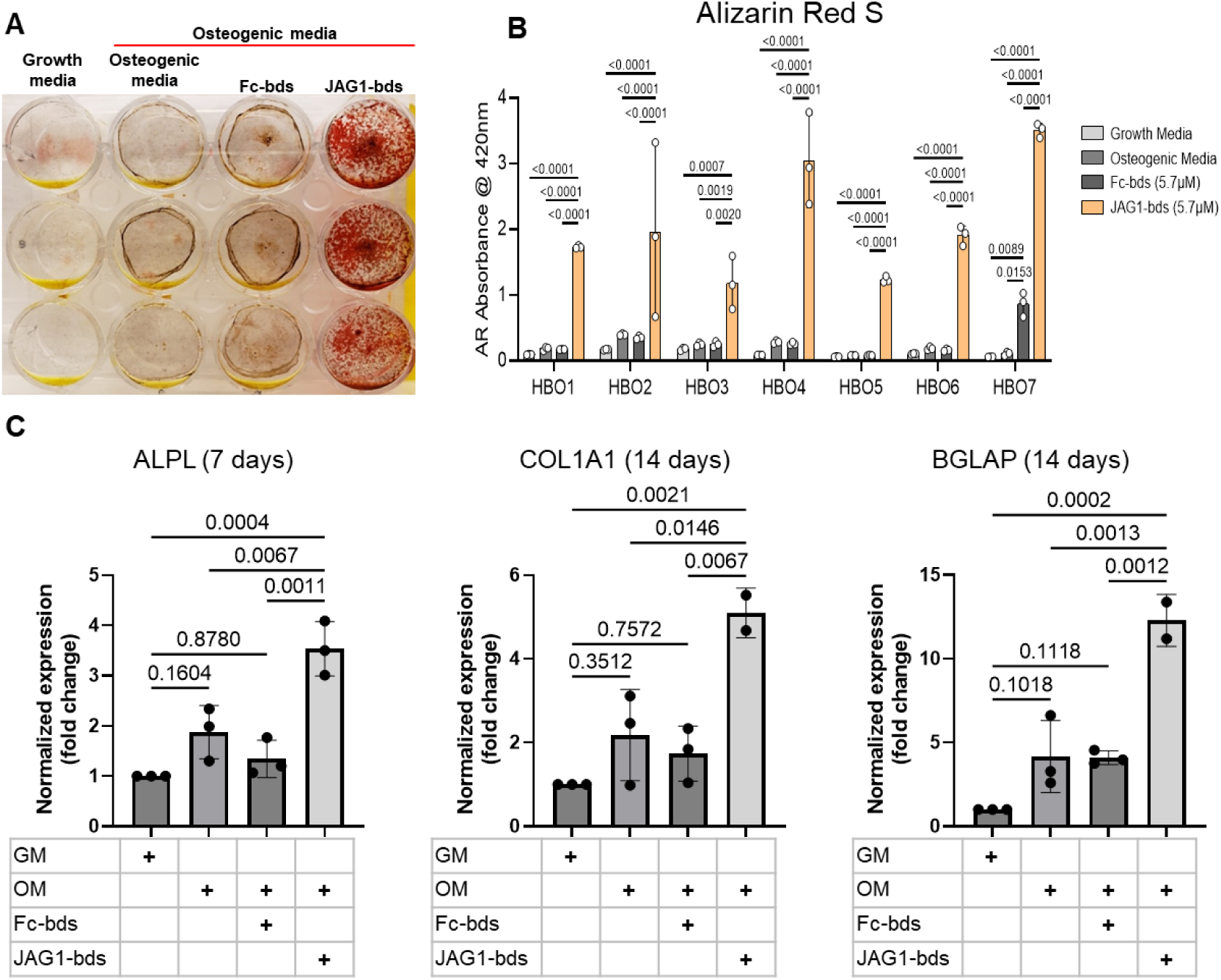
JAGGED1-induced mineralization and gene expression in HBO cells: Seven HBO cell lines were treated with growth media alone, osteogenic media alone or with Fc-Dynabeads (5.7 μM) or JAG1-Dynabeads (5.7 μM). The cells were half-fed every 5 days. On day 21 cells were fixed with 50% ethanol and thereafter, stained with 1% Alizarin Red S. (B) Alizarin Red S dye was extracted from Alizarin Red S-stained cells using a 1:10 dilution of acetic acid and water, and the absorbance was read at 420 nm. Representative image of HBO2. Data represents the mean values of three technical replicates per cell line (mean ± SD, one way ANOVA with Tukey post-hoc). (C) HBO1 primary cell line was grown in triplicate and treated with growth media alone, osteogenic media alone or with Fc-Dynabeads (5.7 μM) or JAG1-Dynabeads (5.7 μM). The cells were half-fed every 5 days and collected at 7 days, 14 days, and 21 days. qRT-PCR was performed (see Methods). Data was normalized to growth media with GAPDH as the reference gene. Data represents the mean values of three biological and two technical replicates per condition (mean ± SD, ordinary one-way ANOVA with Šídák’s multiple comparisons test, with single pooled variance).

### Delivery of JAG1-Dynabead-PEG-4MAL with pediatric HBO cells repairs critically-sized cranial defects

Since we observed an induction of osteoblast commitment and differentiation in pediatric HBO cells in vitro, we next assessed whether co-delivery of JAG1-presenting hydrogels with pediatric HBO cells act similarly in murine craniofacial defects. We recently reported that JAG1 can induce murine CNC-cell osteoblast commitment and repair cranial bone defects in vivo (37). To establish a more translatable use of JAG1 in treating CF bone loss, we assessed whether JAG1 can stimulate human cells (pediatric HBO) to facilitate bone regeneration in NOD-SCID mice, to prevent graft rejection of the HBO cells. JAG1-Dynabead-PEG-4MAL hydrogels also encapsulating pediatric HBO cells obtained from 3 separate donors were implanted in critically-sized parietal bone defects (4 mm) in NOD-SCID mice (n = 4 – 6 per donor, 13 – 15 total) (**Supplementary Figure 3**). The volume of bone regenerated by the JAG1-PEG-4MAL-pediatric HBO hydrogel with or without DAPT, an inhibitor of NOTCH canonical signaling, was measured using μCT, and compared to HBO cells alone, Fc-bds (20 μM) + BMP2 (2.5 μM) treatments. To maintain continuous osteogenic induction, additional JAG1-PEG-4MAL-pediatric HBO cells and control hydrogels were injected into the defects transcutaneously at week four. After eight weeks, we quantified differences in bone volume (BV) within the cranial defect using μCT analysis (**Figure 2A & 2B)**. We observed that there was minimal bone regenerated in mice treated with cells alone. As expected, BMP2 significantly increased regenerated bone volume (p = 0.0002) compared to the cells alone group. The bone volume regenerated by JAG1-bds in the absence (fold change: 1.5) and presence of DAPT (fold change:1.6) was significantly higher compared to the cells alone treatment group (p = 0.0092, 0.0021 respectively). There was no sex-based difference in regenerated bone volume. An initial pilot study also demonstrated no difference in bone regeneration between an Empty Defect model (no HBO cells) and a Cells Alone group (**Supplementary Figure 4**). In **Figure 2C**, paraffin sections of mouse skulls were stained with Masson trichrome stain, as described in methods under histology. Qualitative assessments of the stained sections revealed increased collagen (blue color) in samples obtained from mice treated with BMP2, as expected, and with JAG1-bds and JAG1-bds + DAPT compared to the mice treated with cells alone. To validate the results observed in the samples stained with Masson trichrome, we performed immunohistochemical DAB staining with additional sections from the same mice using rabbit IgG monoclonal anti-COL1A1 antibody. These results corroborate what was revealed with the Masson trichrome staining, that mice treated with cells alone had less collagen production (**Supplementary Figure 5**). This suggests that JAG1 can be used as a bone regenerative therapy where JAG1 induces bone regeneration independently of NOTCH canonical signaling in human cells.

**Figure 2:**
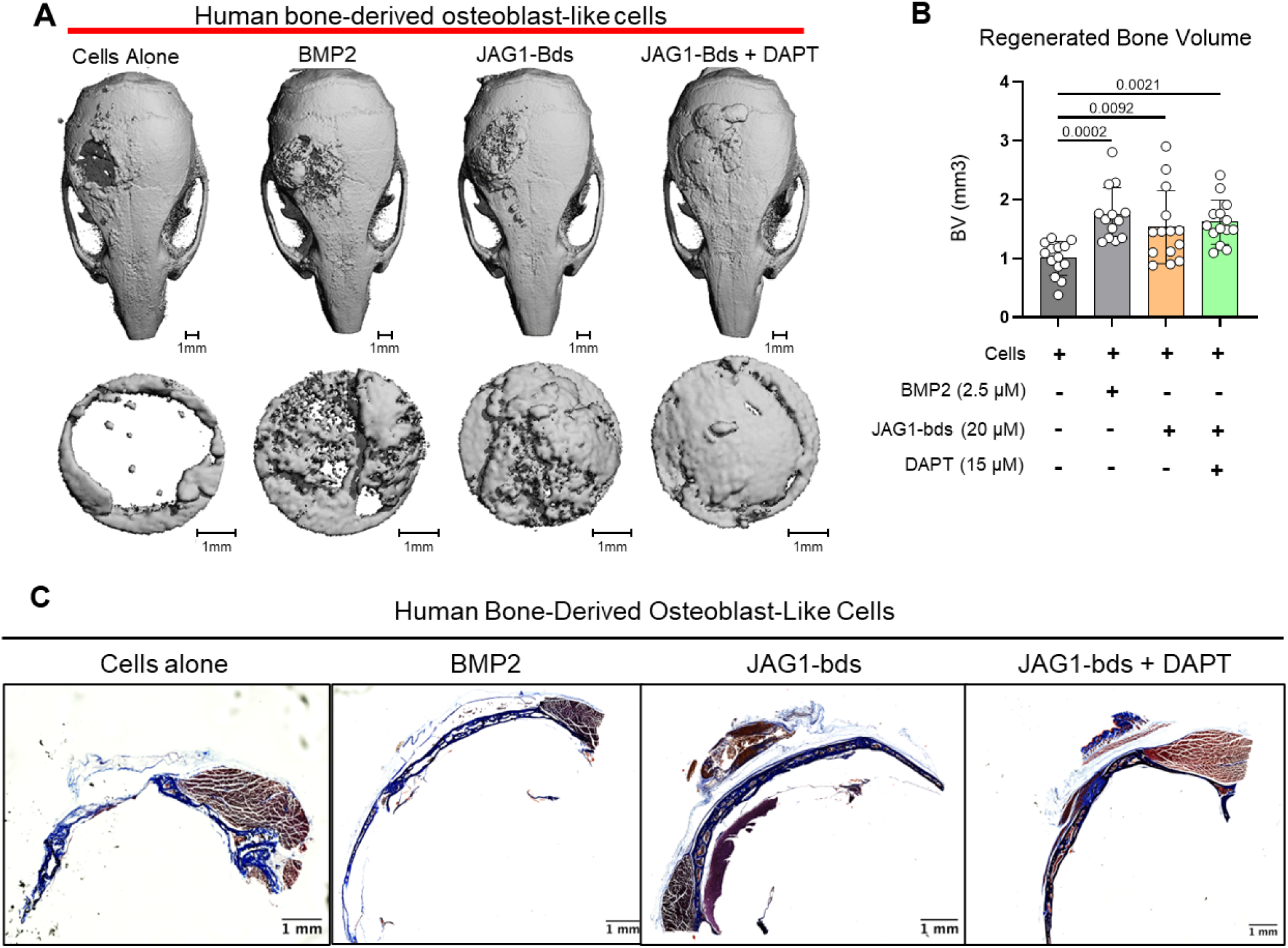
JAG1 delivery in a PEG hydrogel stimulates bone regeneration in a critical-sized bone defect mouse model: HBO cells alone or in the presence of JAG1-Dynabeads complex (20 μM) ± DAPT and BMP2 (2.5 µM)+ Fc-Dynabeads were incorporated in 4% PEG-MAL hydrogels and implanted into 4 mm critical-sized defects in the parietal bones of 6-8-week old NOD SCID mice (n = 4 – 6 per HBO-cell donor, 13 – 15 total) as two separate doses (Initial dose, Week four). After eight weeks, we quantified differences in regenerated bone volume within the defect and compared them between experimental groups by μCT analysis. (A) μCT reconstructions of defects. (B) Quantification of regenerated bone volume. Data are presented as mean (n = 13-15) ± SD with p-values reported (one way ANOVA with Šídák’s multiple comparisons test). (C) shows representative sections of the defect area on skulls from mice from all experimental groups stained with Masson Trichrome stain.

### Transcriptional profiling of cultured HBO cells reveals genes regulated by the non-canonical NOTCH pathway

We previously found that JAG1 can activate a NOTCH non-canonical JAK2-STAT5 signaling pathway in mouse CNC cells, which stimulated expression of osteoblast genes (*Runx2* and *Bglap* (Osteocalcin)) as well as osteoblast commitment and proliferation (1, 37). Thus, we asked if JAG1 would have similar effects for a non-canonical NOTCH pathway on HBO cells. To evaluate this, we cultured HBO cells from a single donor in triplicate and conditioned for 24 hours with vehicle, DAPT, JAG1-bds, or both. Comparison of the JAG1-bds and JAG1- bds+DAPT conditions revealed clusters of genes that were up- or down-regulated by JAG1 stimulation and remained with inhibition of NOTCH, i.e., defining the non-canonical pathway (**Figure 3A**). Analysis of differentially expressed genes (DEGs) in JAG1-bds and JAG1-bds+DAPT groups compared to no treatment revealed a total of 448 up-regulated genes and 435 down-regulated genes in the non-canonical pathway (**Figure 3B**, **Supplementary Table 2**). These include up-regulation of genes involved in osteoblast commitment (*RUNX2*), matrix remodeling (*MMP3*), and diverse cytokines and chemokines (*CCL5, CXCL1, CXCL6*) as part of the non-canonical pathway (**Figure 3C**). Gene ontology analysis of the up-regulated genes in the non-canonical pathway revealed significant over-representation of GO Terms associated with *RUNX2*, cytokine signaling, *NF-κB*, and cell cycle (**Figure 3D**). More interestingly, the PIP3 activating *AKT* signaling pathway was upregulated by JAG1-bds treatment, suggesting that JAG1 can activate NOTCH non-canonical signals via the AKT pathway. Collectively, these data suggest that JAG1 has a profound NOTCH non-canonical effect on HBO cells that stimulates HBO cell-osteoblast commitment and differentiation leading to HBO-cell induced bone formation.

**Figure 3:**
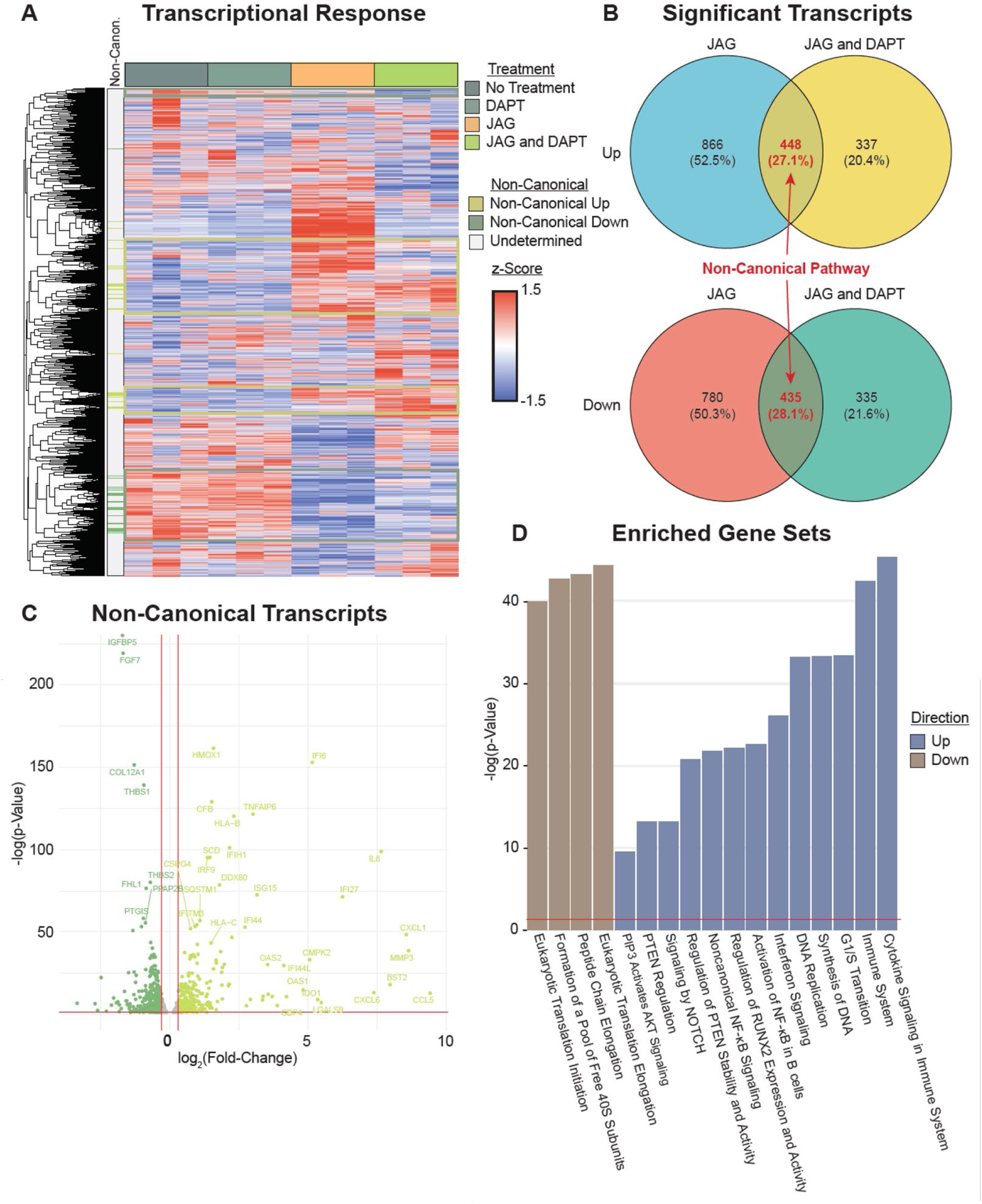
Transcriptional profiling reveals genes and pathways stimulated by non-canonical NOTCH signaling. (A) RNAseq reveals clusters of genes associated with the non-canonical NOTCH pathway (rows are z-scored, side color bar identifies up- and down-regulated DEGs stimulated by the non-canonical pathway). (B) Overlapping DEGs in the JAG1-bds vs no-treatment and JAG1-bds + DAPT vs no-treatment comparisons reveal the non-canonical pathway (DEseq2). (C) Overlapping DEGs from JAG1-bds + DAPT vs no-treatment comparison. (D) Gene ontology over-representation test reveals significantly enriched up- and down-regulated pathways (FDR adjusted).

### JAG1 induces increased cytokine production and phosphorylation of NOTCH non-canonical pathway targets in pediatric HBO cells

Having observed that JAG1 induced osteoblast commitment of murine CNC cells via a NOTCH non-canonical pathway (JAG1-JAK2), we sought to determine whether JAG1 activated NOTCH non-canonical pathways and targets in the pediatric HBO cells. Serum-starved pediatric HBO cells were subsequently treated with JAG1-bds (5.7 μM) with or without DAPT (15 μM), to block the NOTCH canonical pathway, as a time course stimulation for 5, 10, 15, and 30 minutes. Phosphorylation levels for signaling molecules were assessed via Luminex-based multiplex assays. We observed significantly increased phosphorylation of multiple signaling molecules, including STAT5, AKT, P38, JNK, NF-κB, and p70 S6K in JAG1- bds-treated cells, even in the presence of DAPT (**Figure 4**).

**Figure 4:**
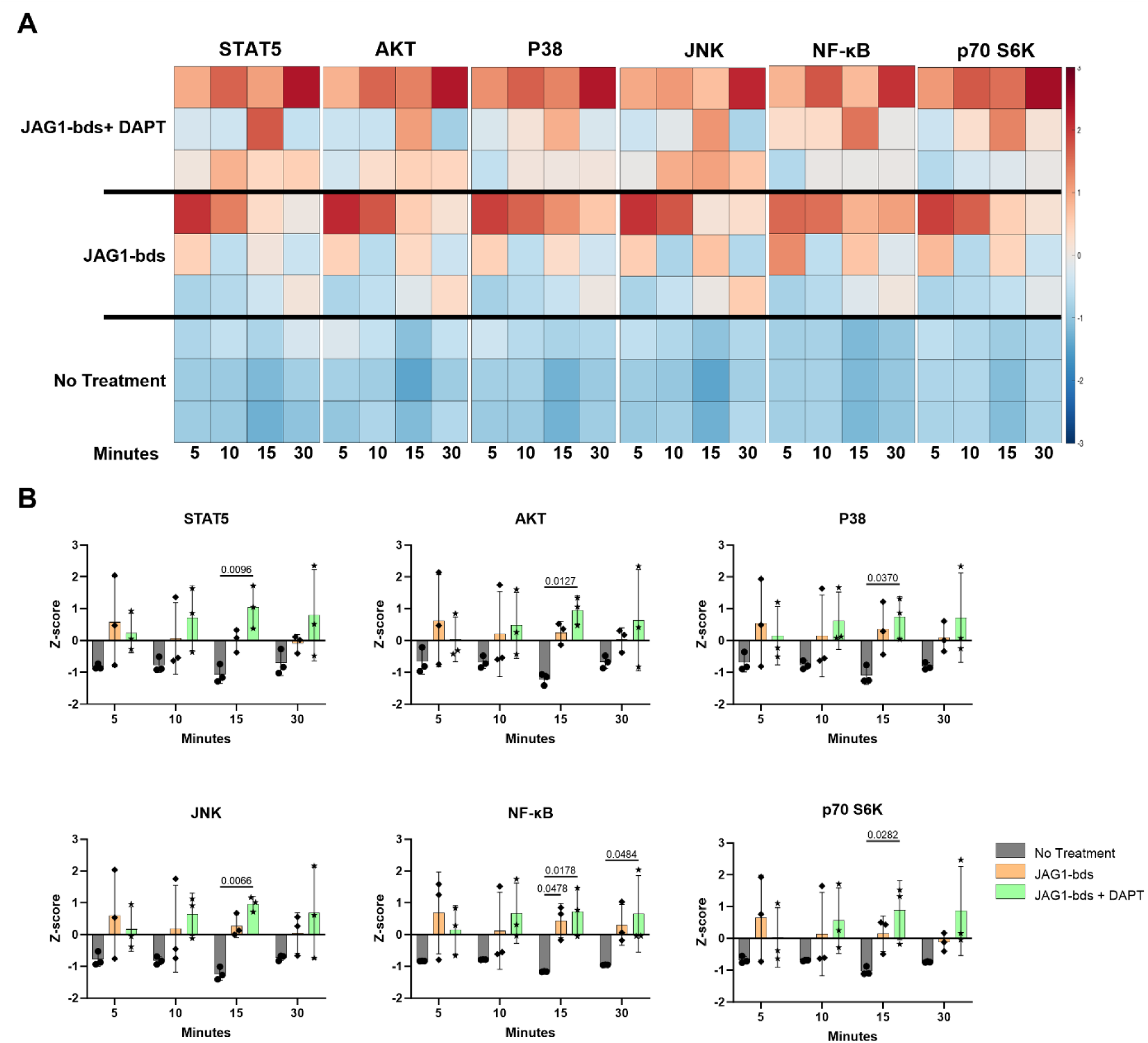
JAGGED1 induces a non-canonical NOTCH pathway in HBO cells. HBO cells undergo mineralization through a non-canonical pathway. Luminex analysis of lysates obtained from three HBO cell lines untreated or treated with Dynabeads-bound recombinant JAG1-Fc fragment (5.7 μM) ± DAPT (15 μM), a NOTCH canonical pathway inhibitor in a time course manner (5, 10, 15 and 30 minutes), A) Heatmaps and B) Z-scores plotted on graphs. Each data point represents mean n = 3 ± SD per cell line with p-values reported.

**Figure 5:**
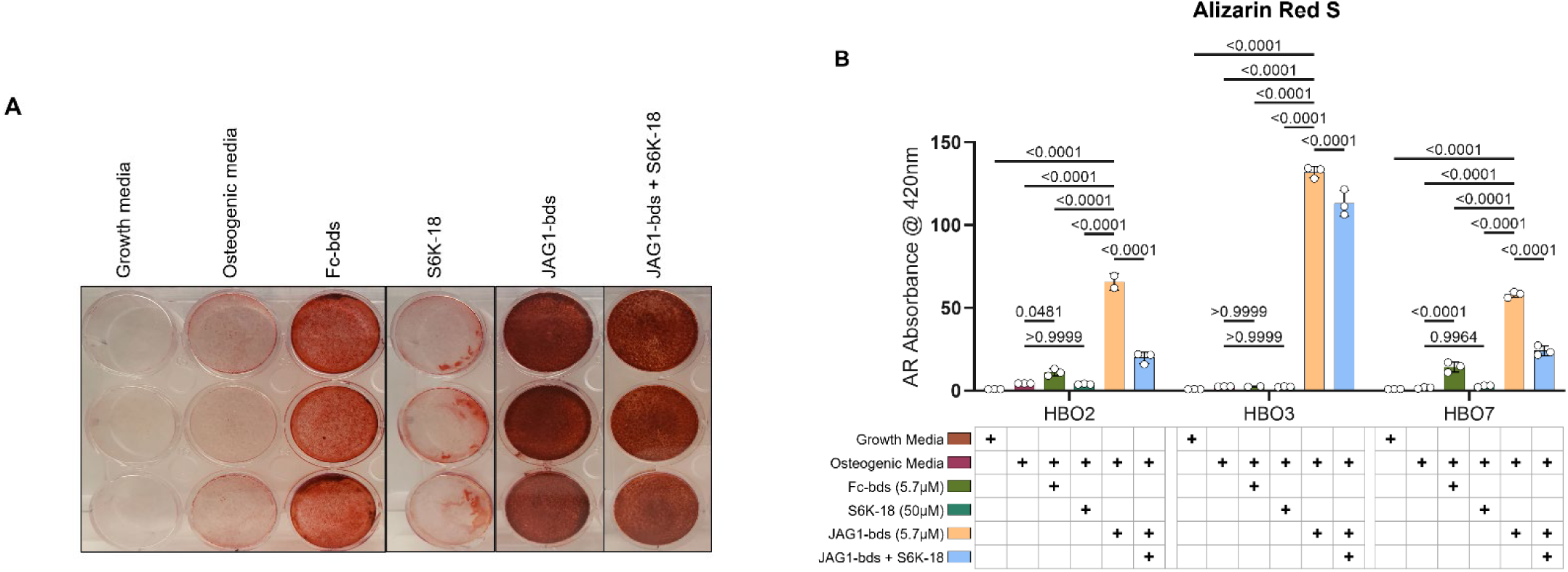
p70 S6K is an essential target during JAGGED1-induced mineralization of HBO cells: HBO cells were treated with growth media alone, osteogenic media alone or with Fc-Dynabeads (5.7 μM), S6K-18 alone (a p70 S6K phosphorylation inhibitor) (50 μM), and JAG1-Dynabeads (5.7 μM) alone or in combination with S6K-18 (50 μM). The cells were half-fed every 5 days. On day 21 cells are fixed with 50% ethanol and thereafter, stained with 1% Alizarin Red S. (B) Alizarin Red S dye was extracted from stained cells using a 1:10 dilution of acetic acid and water, and the absorbance was read at 420nm. Representative image of HBO7. Data represents mean n = 3 ± SD per cell line with p-values indicated.

We have previously shown that JAG1 induced the phosphorylation of STAT5 during CNC cell differentiation to osteoblasts (37). Prior studies emphasize the importance of non-canonical signaling, crosstalk between NOTCH, and other cellular signaling mechanisms. For example, the WNT pathway cross-talks with the NOTCH canonical pathway during vascular morphogenesis (38). Furthermore, previous reports have shown that STAT5 is essential for AKT-p70 S6K activity during lymphocyte proliferation in patients with leukemias and lymphomas (36). Similarly, the P38 pathway has been shown to activate the mTOR-p70 S6K pathway during oxidative stress in mouse embryonic fibroblast cells which culminates in upregulation of antioxidative enzymes that assist in reactive oxygen species removal and thereafter increase cell survival (39).

Also shown previously, JNK phosphorylates p70 S6K to induce osteoblast proliferation and differentiation of MC3T3 cells, and can crosstalk in other physiological systems, for example, during hepatocyte proliferation (30–32). A study by Miwa et al, in 2012, showed that AKT-mTOR-p70 S6K, ERK, and NF-κB were involved together in proliferation of osteosarcoma cells and these pathways could be inhibited by caffeine thereby decreasing tumor burden (40). We observe in our results that many of the pathways activated by JAG1 in the pediatric HBO cells lead to the phosphorylation of p70 S6K, downstream. These findings implicating phospho-protein signaling pathways, especially NF-kB and AKT (downstream of PTEN) are consistent with our transcriptional profiling (**Figure 3**) and collectively implicate p70 S6K as a potential downstream contributor to JAG1-induced, pediatric HBO cell-mediated bone regeneration. Therefore, we proceeded to determine if p70 S6K is an essential downstream target of the JAG1-induced NOTCH non-canonical signaling in pediatric HBO cells (**Figure 5**).

### Inhibition of phosphorylated p70 S6K leads to inhibition of JAG1-induced mineralization of pediatric HBO cells

As shown previously, we measured significantly increased alkaline phosphatase production as well as mineralization in JAG1-bds treated samples compared to all other treatments. We also showed an increase in the phosphorylation of p70 S6K. According to multiple reports in literature, phosphorylation of p70 S6K occurs downstream of multiple pathways (AKT, JAK-STAT, and P38) that we identified in JAG1-induced pediatric HBO cells (36, 39, 40). Thus, we next tested whether the phosphorylation of p70 S6K is an essential downstream target of JAG1-NOTCH during HBO cell mineralization. HBO cells were treated with growth media alone, osteogenic media alone, or Fc-bds (5.7 μM) as negative controls, and JAG1-bds (5.7 μM) with or without S6K- 18, an inhibitor of phosphorylated p70 S6K (41), in the presence of osteogenic media. On day 21, the cells were stained for mineralization using Alizarin Red S stain. As shown in **Figure 5**, we observed that negative controls did not induce mineralization of the HBO cells while JAG1-bds induced significantly higher levels of mineralization, as expected. S6K-18-treated samples showed partial inhibition of 50.2% of the JAG1-bds- induced mineralization (p = 0.0015) and not 100%, possibly because the NOTCH canonical pathway was not inhibited in these samples. Inhibition of JAG1-bds-induced mineralization by S6K-18 treatment alone was significant compared to mineralization induced by JAG1-bds alone suggesting that the phosphorylation of p70 S6K is an essential event downstream of JAG1-NOTCH in JAG1-stimulated HBO cells. Additionally, inhibition of JAG1-bds-induced mineralization was observed with DAPT, an inhibitor of canonical NOTCH signaling (**Supplementary Figure 6**). Subsequent repeat experiments involved collecting the cells on days 9, 14, and 21 for qRT-PCR analysis. While inhibition of NOTCH and p70 S6K decreased mineralization in our mineralization assay, there are no statistically significant changes in gene expression for *ALPL*, *COL1A1*, or *BGLAP* (**Supplementary Figure 7).** These results suggest that the HBO cells phenotypes are maturing into osteocytes and that inhibiting p70 S6K hinders the cellular ability to mineralize but not the cell phenotype progression.

## 4. Discussion

We and others have previously demonstrated that JAG1 exhibits osteo-inductive properties in murine cell lines (1, 37, 42–44). JAG1 also induces the expression of pre-osteoblast genes like *Runx2* in murine CNC cells, even when the canonical NOTCH pathway is disabled using DAPT (1, 28), and further promotes CNC cell commitment to osteoblast differentiation via a JAG1-JAK2 non-canonical signaling pathway (1, 43). Thus, we hypothesized that JAG1 can induce osteoblast differentiation and mineralization of pediatric human bone osteoblast-like (HBO) cells, and the delivery of JAG1 with pediatric HBO cells constitutes an effective treatment for inducing bone regeneration in a pediatric craniofacial bone loss model.

To enhance the clinical translatability of this study, our lab derived human bone-derived osteoblast-like (HBO) cells by collagenase digestion of human pediatric fibula bones. JAG1 has previously been shown to induce survival and proliferation of human mesenchymal stem cells, which are osteoblast precursors and various other human cell types, for example LNCaP which is a prostate cancer cell line, glioma cells, and intestinal epithelial cells during progression of colorectal cancer (45, 46). Osathanon et al, showed that tissue culture plate surface-immobilized JAG1 stimulated osteoblast proliferation and differentiation in iliac bone-derived cells (47). As seen in **Figure 1A &1B**, we also observed increased mineralization of the JAG1-bds-treated HBO cells compared to other controls (growth media alone, osteogenic media alone, and Fc-bds) in HBO cell lines obtained from seven different pediatric fibular samples. The ability of Fc-bds to stimulate mineralization in vitro is previously described in the literature but has not been found to be osteogenic in vivo (1, 37). This suggests that JAG1 reliably induces the human osteoblast-like primary cell expansion and differentiation in multiple human bone cell lines. These results suggest that HBO cells behave similarly to murine CNC cells in vitro and applying these humanized experiments to critically-sized defects will assess JAG1 as a potential bone regenerative therapeutic.

To confirm that JAG1 induces pediatric HBO cells to facilitate osteogenesis in vivo, as it did in vitro, we tested this strategy using an in vivo model of repair using murine craniofacial defects. As shown in **Supplementary Figure 3A & 3B**, PEG-4MAL hydrogels encapsulating JAG1-Dynabeads and pediatric HBO cells were implanted in critically-sized parietal bone defects (4 mm) in athymic nude mice (NOD-SCID), which were used to prevent rejection of human cells. Four weeks later the mice received a second dose of treatments (**Supplementary Figure 3C**) via a transcutaneous injection to maintain the osteogenic signaling. Intermittent treatment with bone-regenerative therapeutics, like Parathyroid hormone (1–34), (48, 49) has been shown to lead to anabolic increase in bone mineral density. The ORTHOUNION clinical trial which aimed at enhancing bone healing in long bone nonunion fractures focused on testing the treatment involving two different sequential doses of expanded bone marrow-derived mesenchymal stem cells (50). In our study, eight weeks after the initial dose, the regenerated bone volume was measured using µCT. As seen in **Figure 2B**, the fold change of bone volume regenerated by JAG1-bds in the presence of DAPT (fold change:1.6, normalized to cells alone treatment) was comparable to JAG1-bds alone treatment (fold change: 1.5, p = 0.9583), and was significantly higher compared to the cells alone treatment group (p = 0.0038). The inhibition of NOTCH canonical signaling in mesenchymal stem cells using DAPT has been shown previously to enhance osteogenesis (51). A rheumatoid Arthritis C57BL/6 SCID mouse model carrying a human *TNF* transgene when treated with intermittent doses of DAPT showed improved bone regeneration (52). This supports our current results that JAG1 induces bone regeneration independently of NOTCH canonical signaling in the HBO cells. However, structural and mechanical properties of the bone will require characterization in the future using biomechanical testing.

As shown in our previous publications, JAG1 induced osteoblast commitment of murine CNC cells via a NOTCH non-canonical pathway. In our current data, it is observed that JAG1 induced HBO cells to regenerate bone and repair cranial defects even in the presence of a NOTCH canonical pathway inhibitor (DAPT), which suggests that JAG1-induced pediatric HBO cell-facilitated bone regeneration occurs via a NOTCH non-canonical pathway. Therefore, to identify the NOTCH non-canonical pathway targets that are activated by JAG1-bds in HBO cells, we isolated RNA from the JAG1-bds-treated HBO cells, subjected it to RNA sequencing and performed pathway analyses on the data obtained. The data showed that a total of 448 genes were up-regulated, and 435 genes were down-regulated in JAG1-bds and JAG1-bds+DAPT groups compared to no treatment and thus these genes participate in the non-canonical NOTCH signaling pathway (**Figure 3B**, **Supplementary Table 2**). Some of the genes that are upregulated as part of the non-canonical NOTCH pathway are known to be involved in osteoblast commitment (*RUNX2*), matrix remodeling (*MMP3*), and diverse cytokines and chemokines (*CCL5* and *CXCL1*). RUNX2 is commonly known to be expressed by osteoblasts as a sign of their recruitment into the lineage (53). MMP3 is also abundantly expressed by osteoblasts and is an important regulator of bone remodeling. MMP3 is an enzyme that is essential in the processing of collagen on bone surface, which in turn is necessary for osteoclast recruitment and bone resorption (54). Chemokine CCL5 is abundantly expressed by osteoblasts, and it is also involved in recruitment of osteoblast progenitor cells (55). PTH/PTH1R induces the differentiation of osteoblasts, and it has been shown that CXCL1 serves as an intermediate during this process (56). Gene ontology analysis of the up-regulated genes in the non-canonical pathway revealed significant enrichment of GO Terms associated with *RUNX2*, cytokine signaling, and cell cycle as described earlier are involved in osteoblastic cell proliferation, recruitment, and differentiation leading to bone remodeling. *NF-κB* was also upregulated and it is known to be a radiation-induced pro-survival factor in human osteoblastic cells (57). More interestingly, the PIP3 activating AKT signaling pathway was upregulated by JAG1-bds treatment, suggesting that JAG1 can activate NOTCH non-canonical signals via the AKT pathway. Collectively, these data reveal that JAG1 has a profound NOTCH non-canonical effect on HBO cells that stimulates HBO cell-osteoblast commitment and differentiation, and HBO-cell induced bone formation.

We also obtained lysates from HBO cells treated with JAG1-bds in the presence and absence of DAPT and subjected them to Luminex-based multiplex assays. The Luminex-based assays demonstrated that JAG1 induces the phosphorylation of various signaling targets, including STAT5, AKT, P38, JNK, NF-κB, and p70 S6K, between 15 to 30 minutes, as shown in **Figure 4**. As discussed earlier, prior studies emphasize the importance of non-canonical signaling, crosstalk between NOTCH, and other cellular signaling mechanisms (38) These studies show that STAT5 is essential for AKT-p70 S6K activity during lymphocyte proliferation in patients with leukemias and lymphomas (36), and that AKT-mTOR-p70 S6K, ERK, and NF-κB were involved together in proliferation of osteosarcoma cells (40). JNK was previously found to phosphorylate p70 S6K to induce osteoblast proliferation and differentiation of MC3T3 cells (58–60), and that the P38 pathway has been shown to activate the mTOR-p70 S6K pathway during oxidative stress in mouse embryonic fibroblasts (39). Taken together, our results and the results of others demonstrate that the p70 S6K pathway may be a node at which osteoblast induction occurs in HBO cells, making it a potential major contributor to bone regeneration caused by JAG1-induced HBO cells. To test this hypothesis, we proceeded to confirm that p70 S6K is an essential target of the JAG1-induced NOTCH non-canonical signaling in HBO cells (**Figure 5**). p70 S6K is an enzyme that phosphorylates the S6 ribosomal protein to initiate protein synthesis which supports growth, proliferation, differentiation and glucose homeostasis of cells (61–63). We found that JAG1-induced mineralization in HBO cells was predominantly (50.2%) inhibited by S6K-18, an inhibitor of the phosphorylation of p70 S6K, recognizing that other non-canonical signaling pathways are also important in bone regeneration (e.g. p38, AKT). However, inhibition of JAG1-induced mineralization caused by S6K-18 treatment alone was significant compared to that induced by JAG1-bds alone (p = 0.015), suggesting that the phosphorylation of p70 S6K is a significant contributor of osteoblast induction in JAG1-stimulated HBO cells. Thus, p70 S6K is an important downstream target of JAG1-NOTCH in JAG1-stimulated HBO cells.

Studying the mechanisms by which JAG1 induces osteoblast commitment and bone formation will enable new treatment avenues that involve the delivery of not only tethered JAG1 but also the JAG1-NOTCH non-canonical signaling intermediates themselves or their activators as powerful treatment options to induce bone regeneration in CF bone loss injuries. High-throughput screening for existent, FDA-approved drugs, compounds, and small molecules can be used to identify pharmacological activators of p70 S6K, as future directions of this study. Additionally, further investigation of NF-κB and the other downstream signaling targets identified in our Luminex-based assays are currently in the planning stages. These findings can provide powerful treatment options to induce bone regeneration in CF bone loss injuries and avoid the limitations with currently available therapies.

## Acknowledgements

Study design: AK, AIT, SA, AJG, LBW, HD, MHR, TC, SC and SLG. Study conduct: AK, SK, MHR. Data collection: AK, BT, SK, MHR, and IM. Data analysis: AK, BT, SK, MHR, AIT, IM. Data interpretation: All authors. Drafting manuscript: AK and SLG. Revising manuscript content: All authors. Approving the final version of manuscript: All authors. We extend our gratitude to Adrianna Westbrook and Katie Liu from the Pediatric Biostatistics Core, Emory University, for their invaluable advice on how best to perform the statistics on the data in this manuscript. AK and SLG take responsibility for the integrity of the data analysis. Research reported in this publication is supported by the National institute of Health, National Institute of Dental and Craniofacial Research (NIDCR) under award number R01DE031271 and National institute of Health, National Institute of Arthritis and Musculoskeletal and Skin Diseases of the National Institutes of Health under Award Numbers DE026762, R01AR062920, and R01AR062368, respectively. The authors declare no conflict of interest.

## Abbreviations

CF: craniofacial.
BMP2: bone morphogenetic protein-2.
FDA: Food and Drug Administration.
JAG1: jagged canonical Notch ligand 1.
CNC: Cranial Neural Crest cells.
JAK2: Janus kinase 2.
STAT5: signal transducer and activator of transcription 5.
HBO: Human Bone-derived Osteoblast-like (HBO) cells.
p70 S6K: RPS6KB1 ribosomal protein S6 kinase B1.
PTH (1-34): Parathyroid Hormone (1-34).
PTH: parathyroid hormone
PTH1R: parathyroid hormone 1 receptor
VEGFA: vascular endothelial growth factor A.
FGF: fibroblast growth factor.
SDF-1: Stromal Cell-Derived Factor-1;
CXCL12: C-X-C motif chemokine ligand 12.
TGFB2: Transforming Growth Factor-Beta 2.
PRP: platelet-rich plasma.
HES1: Hes family bHLH transcription factor 1.
HEY1: Hairy/enhancer-of-split related with YRPW motif protein 1.
MMP3: matrix metallopeptidase 3
RUNX2: Runt-related Transcription Factor 2.
SP7: Sp7 transcription factor
COL1A1: collagen type I alpha 1 chain
BGLAP: bone gamma-carboxyglutamate protein (Osteocalcin)
ALPL: alkaline phosphatase, biomineralization associated.
GAPDH: glyceraldehyde-3-phosphate dehydrogenase
PBS: Phosphate-buffered Saline.
(PEG)-4MAL: Maleimide end-functionalized four-arm Poly (ethylene glycol).
HEPES: 4-(2-hydroxyethyl)-1-piperazineethanesulfonic acid.
NOD-SCID: Non-Obese Diabetic/Severe Combined Immunodeficiency.
μCT: Micro Computed Tomography.
BV: Bone Volume.
DEXA: Dexamethasone.
βGP: Beta – Glycerophosphate.
DAPT: N-[N-(3,5-Difluorophenacetyl)-L-alanyl]-S-phenylglycine t-butyl ester.
ANOVA: Analysis of Variance.
SD: standard deviation.
DMEM: Dulbecco′s Modified Eagle′s Medium.
FBS: fetal bovine serum.
AKT: AKT serine/threonine kinase 1.
P38: p38 mitogen-activated protein kinase.
JNK: c-Jun N-terminal Kinase
MAPK8: mitogen-activated protein kinase 8.
NFKB (NF-κB): Nuclear factor kappa B protein complex.
WNT: Wingless-related integration site signaling proteins (WNT1-11, WNT16).
mTOR: Mammalian Target of Rapamycin.
MC3T3: Osteoblast precursor cell line derived from Mus musculus (mouse) calvariae.
LNCaP: Cell line derived from a metastatic lymph node lesion of human prostate cancer.
TNF: tumor necrosis factor

---

**Supplementary Figure 1:**
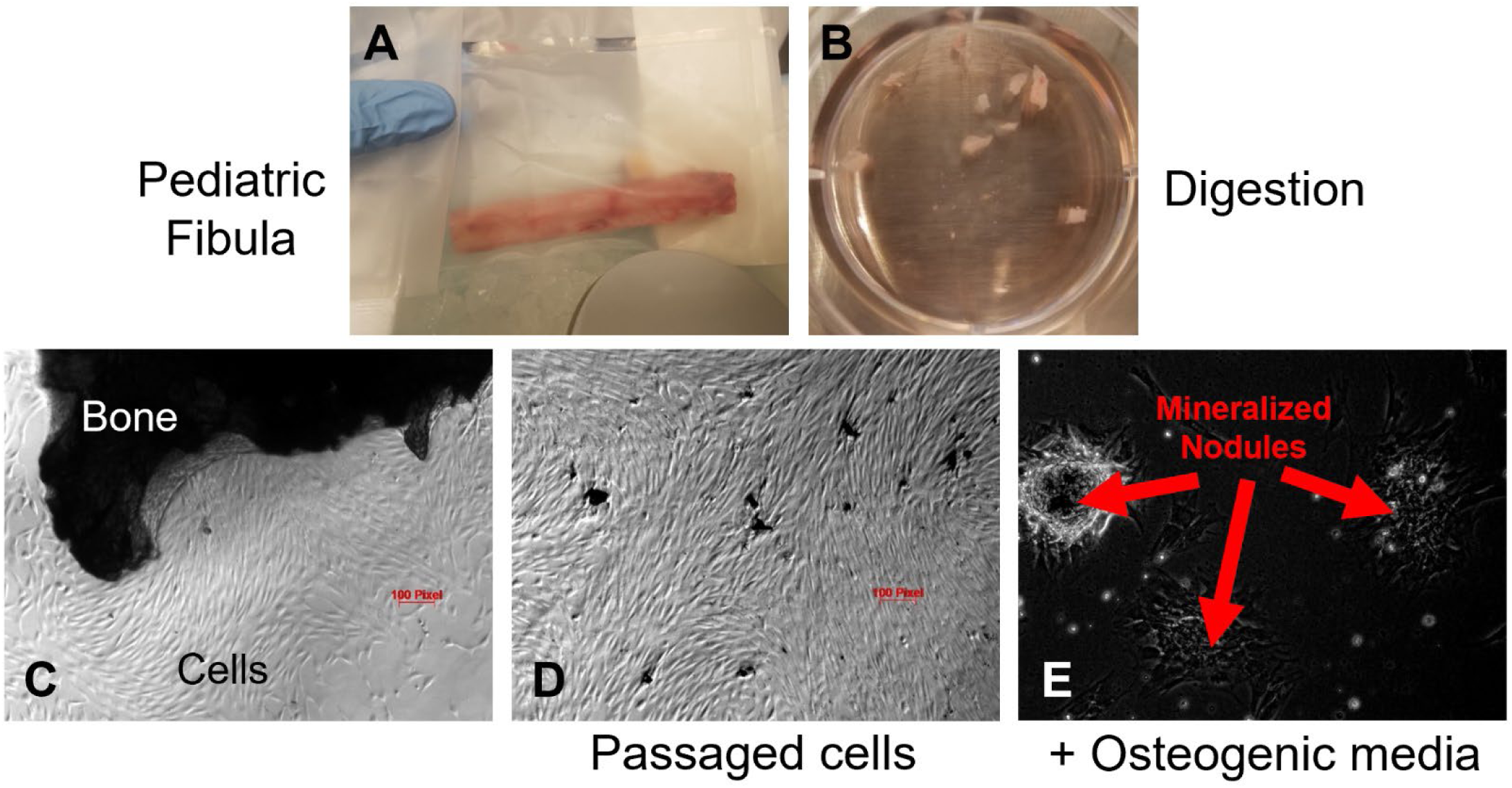
Method of processing human bone samples to produce primary Human Bone-derived Osteoblast-like cell lines. Human Bone-derived Osteoblast-like cells (HBO) were isolated by collagenase digestion of pediatric healthy fibular bones. See Methods.

**Supplementary Figure 2:**
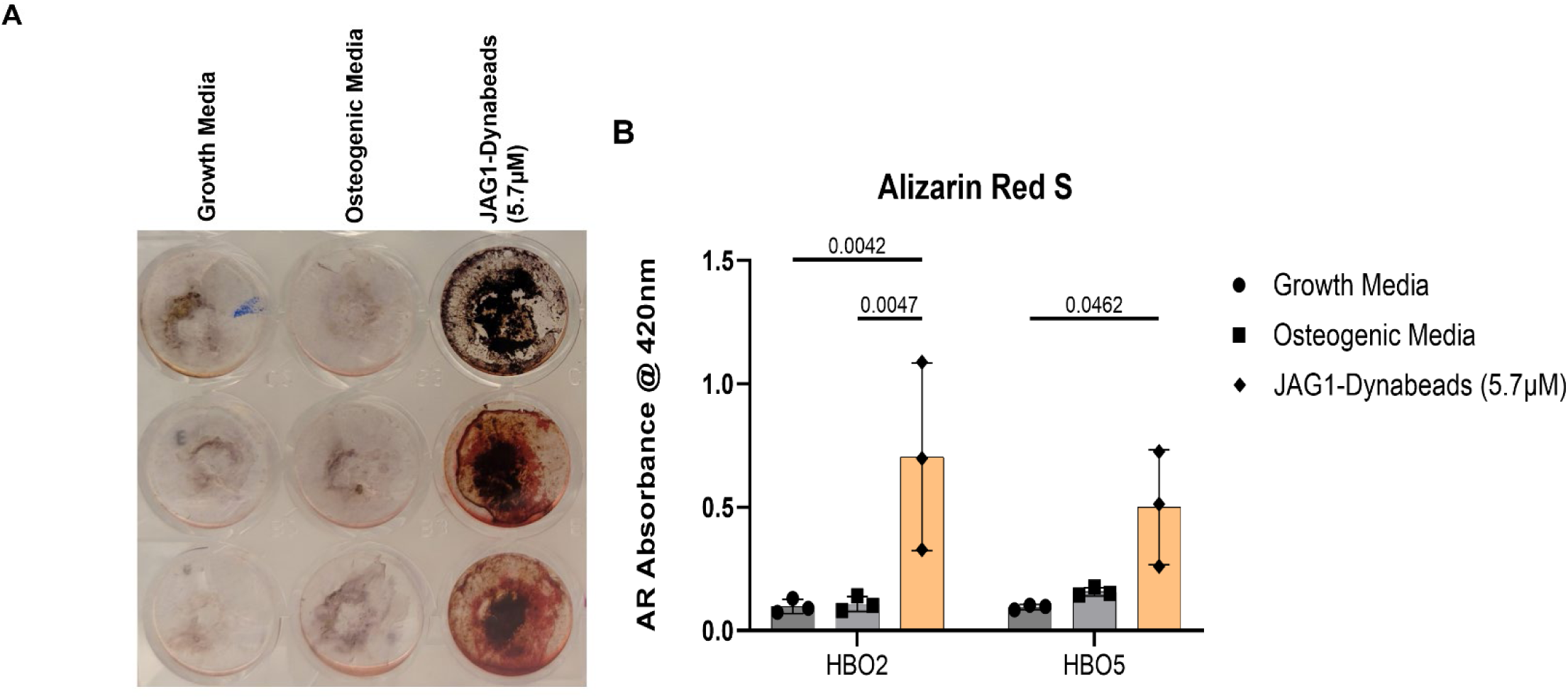
PEG-4MAL hydrogel-encapsulated JAGGED1-induced mineralization of HBO cells. HBO cells alone or in the presence of JAG1-Dynabeads complex (20 μM) were incorporated in 4% PEG-MAL hydrogels and grown in culture. The cells were half-fed every 5 days. On day 21 cells were fixed with 50% ethanol and thereafter, stained with 1% Alizarin Red S. (B) Alizarin Red S dye was extracted from stained cells using a 1:10 dilution of acetic acid and water, and the absorbance was read at 420nm. Data represents mean ± SD per cell line with p-values indicated, n = 3.

**Supplementary Figure 3:**
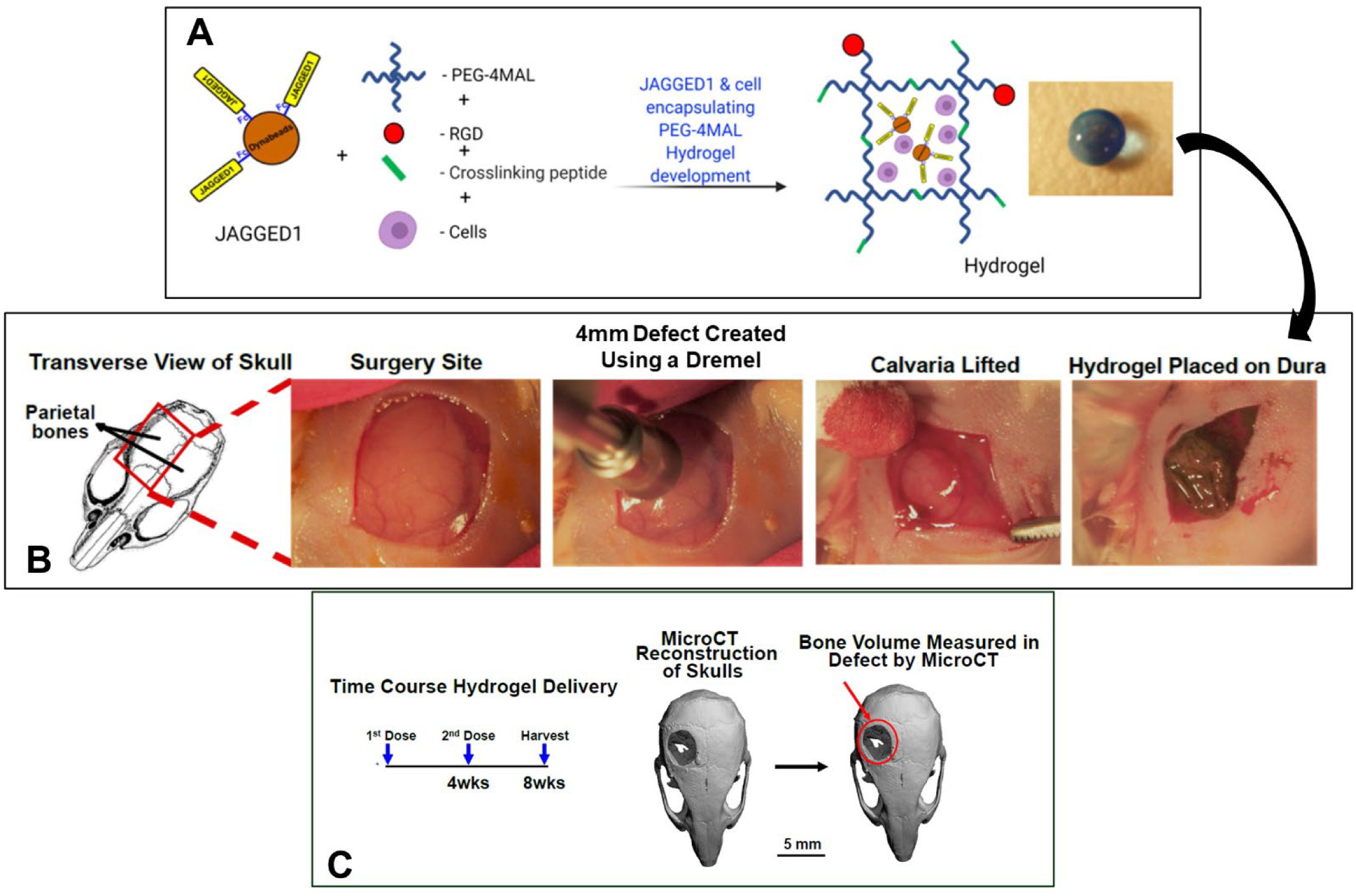
Visual Depiction of Calvarial Defect Studies. HBO cells alone or in the presence of other treatments were incorporated in 4% PEG-MAL hydrogels and implanted into 4 mm critical-sized defects in the parietal bones of 6-8-week-old NOD SCID mice as two separate doses (Initial dose, Week four). After eight weeks, the regenerated bone volume, within the defect and compared them between experimental groups by μCT analysis.

**Supplementary Figure 4:**
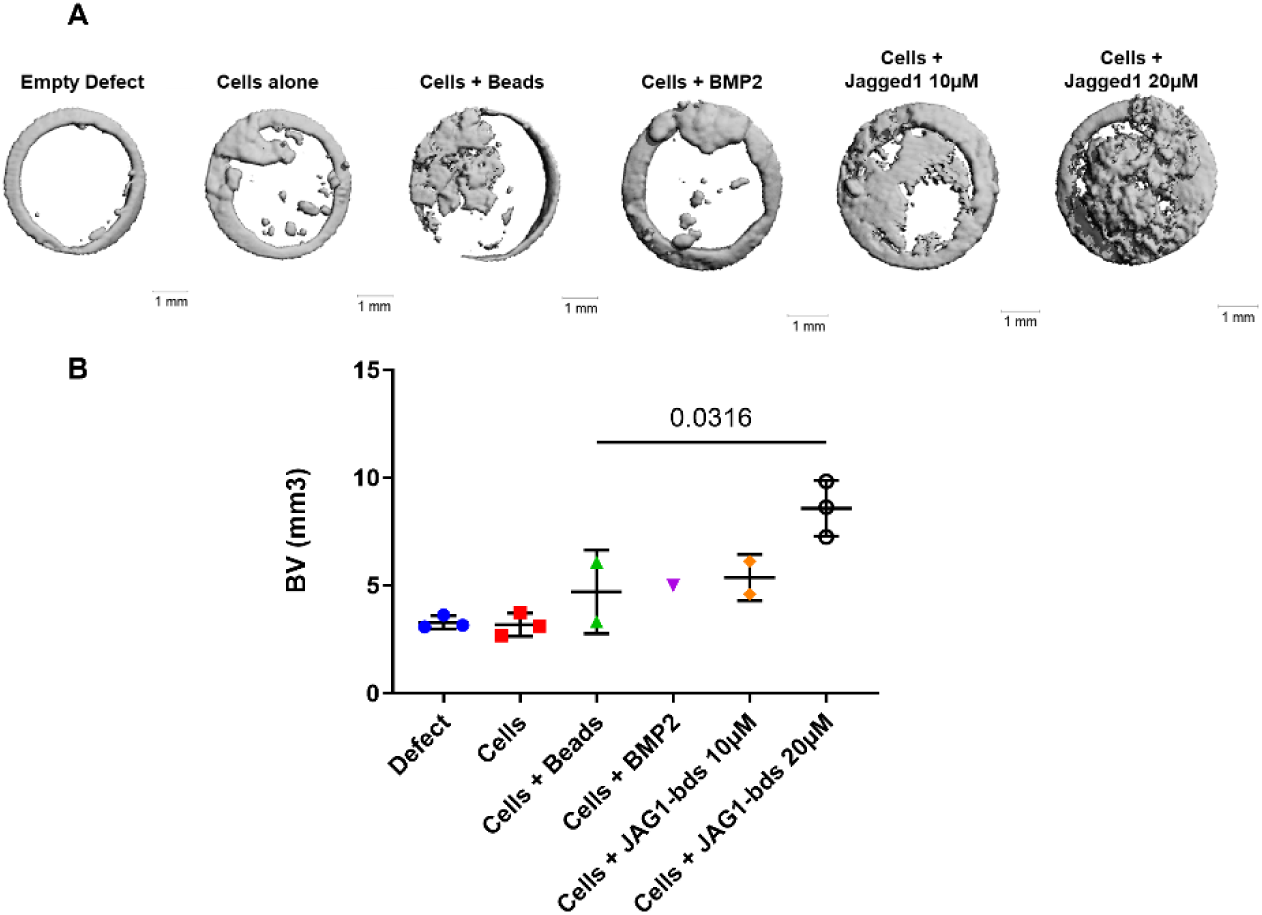
Pilot study of JAG1-bds delivery in a PEG hydrogel stimulates bone regeneration in a critical-sized bone defect mouse model. No cells (Empty Defect) or HBO cells alone or in the presence of JAG1-Dynabeads complex (20 μM) ± DAPT and BMP2 (2.5 µM)+ Fc-Dynabeads were incorporated in 4% PEG-MAL hydrogels and implanted into 4 mm critical-sized defects in the parietal bones of 6-8-week old NOD SCID mice (n = 3) as two separate doses (Initial dose, Week four). After eight weeks, we quantified differences in regenerated bone volume within the defect and compared them between experimental groups by μCT analysis. (A) μCT reconstructions of defects. (B) Quantification of regenerated bone volume. Data are presented as mean (n = 3) ± SD with p-values reported (ordinary one-way ANOVA with Tukey’s multiple comparisons test with a single pooled variance).

**Supplementary Figure 5:**
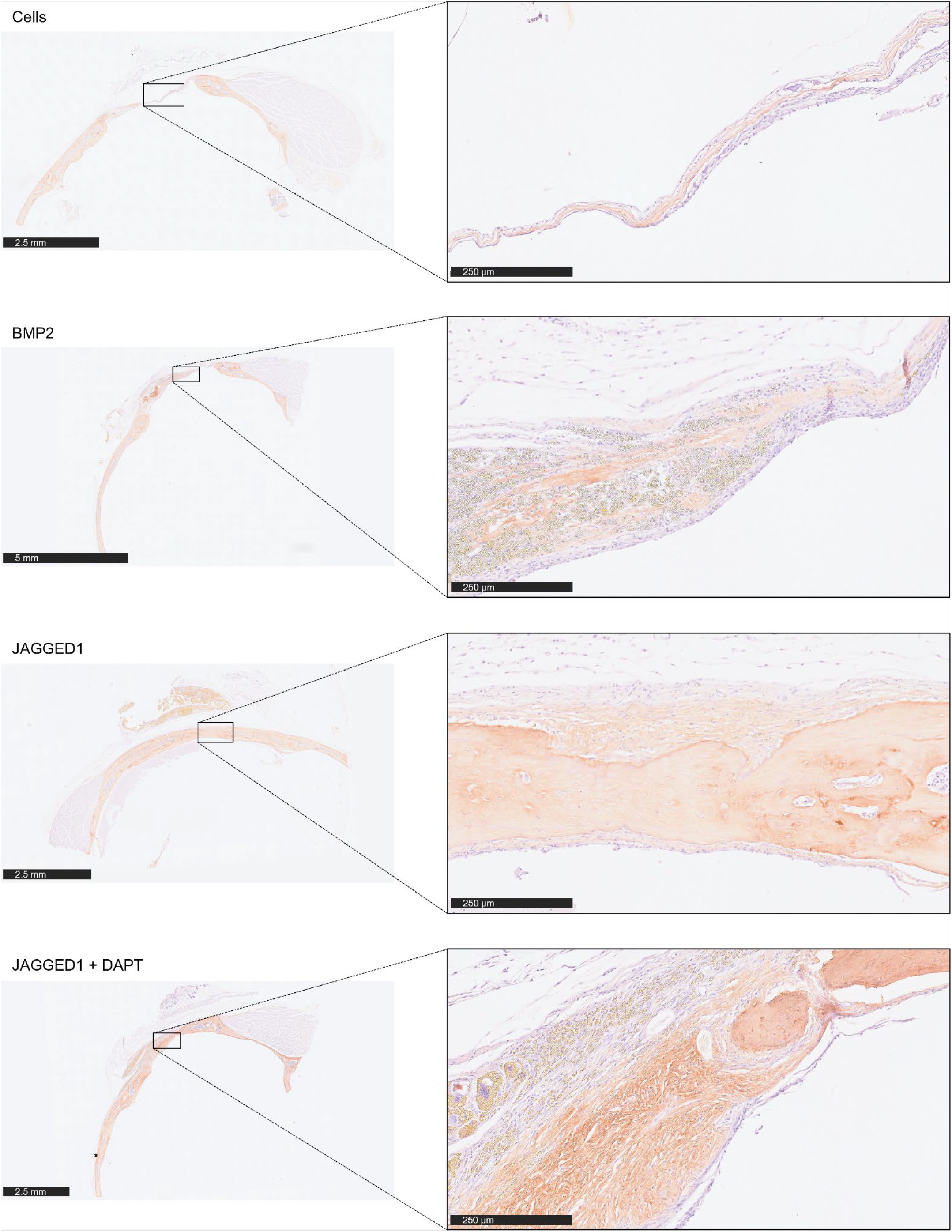
Immunohistochemical staining of calvarial defect tissue for Collagen 1. FFPE tissue from the calvarial defect experiment shown in Figure 2 was sectioned and stained with an antibody for COL1A1 (Cell Signaling #72026S) and counterstained with Hematoxylin. Slides were scanned with the Olympus Nanozoomer whole-slide scanner at 20x.

**Supplementary Figure 6:**
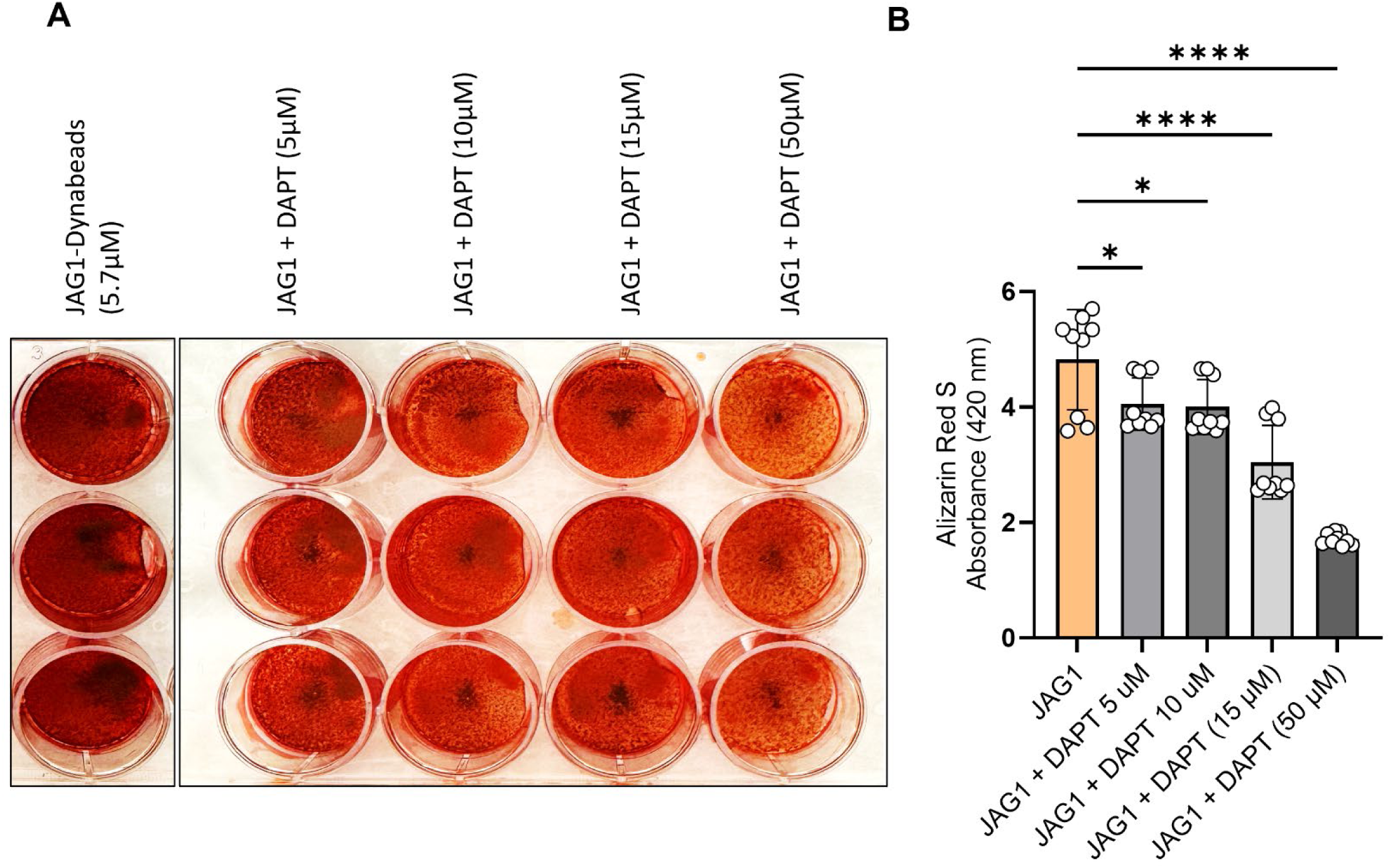
Mineralization assay with DAPT inhibition of the NOTCH canonical pathway: HBO1 cells were treated with JAG1-Dynabeads (5.7 μM) alone or in combination with increasing concentrations of DAPT in a dose response test (50 μM). The cells were half-fed every 5 days. On day 21 cells were fixed with 50% ethanol and thereafter, stained with 1% Alizarin Red S. Representative image of HBO1. (B) Alizarin Red S dye was extracted from stained cells using a 1:10 dilution of acetic acid and water, and the absorbance was read at 420nm. Data represents mean ± SD per cell line with p-values indicated, n = 9.

**Supplementary Figure 7:**
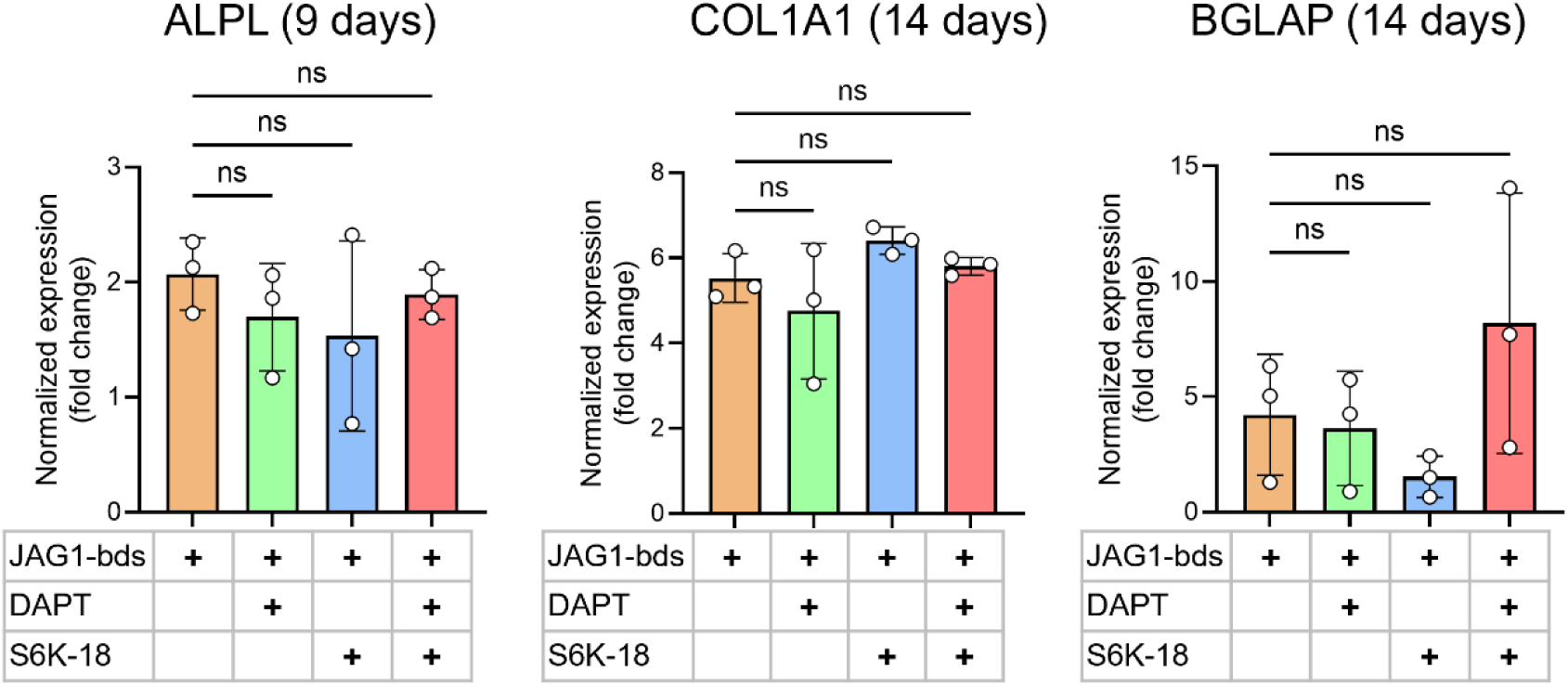
Inhibition of JAGGED1-induced mineralization of HBO cells with inhibitors of NOTCH and p70 S6K: HBO cells were treated with JAG1-bds (5.7 μM) alone or in combination with DAPT (15µM), S6K-18 (a p70 S6K phosphorylation inhibitor) (50 μM) or S6K-18 (50 μM) + DAPT (15 μM). The cells were half-fed every 5 days. Mineralization assays were conducted in triplicate, and cells were collected at days 9, 14, and 21. RNA was subsequently assessed by qRT-PCR as described in Methods. Data was normalized to growth media with GAPDH as the reference gene. These charts compare normalized expression of genes with JAGGED1 stimulation and with inhibitors. Data represents the mean values of three biological and two technical replicates per condition (mean ± SD, ordinary one-way ANOVA with Šídák’s multiple comparisons test, with single pooled variance).

**Supplementary Table 1:** Primer sequences used for qRT-PCR.

**Supplementary Table 2:** List in excel sheet attached shows differentially expressed genes (DEGs) in JAG1 and JAG1+DAPT groups compared to no treatment. Analysis revealed a total of 448 up-regulated genes and 435 down-regulated genes in the non-canonical pathway.

