## Supplemental Table 1 for "Delivery of A Jagged1-PEG-MAL hydrogel with Pediatric Human Bone Cells Regenerates Critically-Sized Craniofacial Bone Defects"

Supplemental Table 1: Primers used in qRT-PCR

| Target | IDT Assay ID | Primer 1 | Primer 2 |
| --- | --- | --- | --- |
| RUNX2 | Hs.PT.56a.19568141 | CTTCACAAATCCTCCCCAAGT | AGGCGGTCAGAGAACAAAC |
| SP7 | Hs.PT.58.26512371 | TCCCTCTCCCTTTTCTCTCTC | GGAGCCATAGTGAACTTCCTC |
| COL1A1 | Hs.PT.58.15517795 | GACATGTTCAGCTTTGTGGAC | TTCTGTACGCAGGTGATTGG |
| BGLAP | Hs.PT.56a.39318706.g | CTCACACTCCTCGCCCTAT | CGCCTGGGTCTCTTCACT |
| Target |  | Forward | Reverse |
| ALPL | Custom | ACA AGC ACT CCC ACT TCA TCT GGA | TCA CGT TGT TCC TGT TCA GCT CGT |
| GAPDH | Custom | AGG GCT GCT TTT AAC TCT GGT | CCC CAC TTG ATT TTG GAG GGA |
| HPRT | Custom | ATT GGT GGA GAT GAT CTC TCT CAA CTT T | GCC AGT GTC AAT TAT ATC TTC CAC AA |
